# Pharmacologic inhibition of lysine specific demethylase-1 (LSD1) as a therapeutic and immune-sensitization strategy in diffuse intrinsic pontine glioma (DIPG)

**DOI:** 10.1101/690966

**Authors:** Cavan P. Bailey, Megan M. Romero, Oren J. Becher, Michelle Monje, Dean A. Lee, Linghua Wang, Joya Chandra

## Abstract

**Background:** Diffuse intrinsic pontine glioma (DIPG) is an incurable pediatric brain tumor. Mutations in the H3 histone tail (H3.1/3.3-K27M) are a feature of DIPG, potentially rendering them therapeutically sensitive to small-molecule inhibition of chromatin modifiers. Pharmacological inhibition of lysine specific demethylase-1 (LSD1) shows promise in pediatric cancers such as Ewing’s sarcoma, but has not been investigated in DIPG, which was the aim of our study.

**Methods:** Patient-derived DIPG cell lines and pediatric high-grade glioma (pHGG) datasets were used to evaluate effects of several LSD1 inhibitors on selective cytotoxicity and immune gene expression. Immune cell cytotoxicity was assessed in DIPG cells treated with LSD1 inhibitors and informatics platforms were used to determine immune infiltration of pHGG and impact on survival.

**Results:** Selective cytotoxicity and an immunogenic gene signature was established in DIPG lines using several clinically-relevant LSD1 inhibitors. Pediatric high-grade glioma patient sequencing data demonstrated survival benefit using this LSD1-dependent gene signature. On-target binding of catalytic LSD1 inhibitors was confirmed in DIPG and pre-treatment of DIPG with these inhibitors increased lysis by natural killer (NK) cells. CIBERSORT analysis of patient data confirmed NK infiltration is beneficial to patient survival while CD8 T-cells are negatively prognostic. Catalytic LSD1 inhibitors are non-perturbing to NK cells while scaffolding LSD1 inhibitors are toxic to NK cells and do not induce the gene signature in DIPG cells.

**Conclusions:** LSD1 inhibition using catalytic inhibitors are both selectively cytotoxic and promote an immune gene signature that is associated with NK cell killing, representing a therapeutic opportunity for pHGG.

**Key points:**

1. LSD1 inhibition using several clinically relevant compounds is selectively cytotoxic in DIPG.
2. An LSD1-controlled gene signature predicts survival in pediatric high-grade glioma patients.
3. LSD1 inhibition enhances NK cell cytotoxicity against DIPG with correlative genetic biomarkers.

**Importance of the study:** This is the first study to evaluate inhibition of LSD1 in a uniformly lethal type of pediatric brain tumor: DIPG. We demonstrate selective cytotoxicity of several clinically relevant compounds against patient derived DIPG cells, and identify an immune gene signature that is upregulated in DIPG cells by catalytic inhibitors of LSD1. This immune gene signature is predictive of prognosis in pHGG, consistent with the rationale of promoting this signature through LSD1 inhibition. NK cell killing of DIPG is enhanced by LSD1 inhibition, providing functional confirmation of this gene signature, and represents the first report of LSD1 inhibition promoting NK cell cytotoxicity of cancer cells. Given the poor prognosis of pHGGs and lack of effective treatments, our results suggest use of LSD1 inhibition as a single agent or in combination with NK cell therapy may be a safe and efficacious strategy.

## Introduction

Pediatric high-grade gliomas (pHGGs) are pathologically diverse yet uniformly highly malignant central nervous system (CNS) cancers, with 5-year survival rates of <10% post-diagnosis. Surgery is often not possible due to tumor locations in the midline and brainstem, which control crucial motor functions such as breathing and heartbeat. Chemotherapy and radiation are the standard of care, but survival benefits are slim with high risks of side effects and decreased quality of life during and after treatment^1^. Immunotherapeutic approaches have had limited success due to the low mutational burden and immunosuppressive microenvironment of pediatric brain tumors, such that adaptive immune interventions including checkpoint blockade are ineffective^2^. Recent efforts to molecularly profile pHGGs have discovered conserved mutations unique to the pediatric age range and anatomical locations^3^. In particular, mutations in histone proteins that package DNA, specifically the alleles H3.3/3.1-K27M and H3.3-G34R/V, confer aberrant gene expression that is thought to drive early development of these tumors in multipotent CNS cells^4^. As such, the World Health Organization (WHO) now recognizes these K27M tumors as separate entities in the pediatric glioma classification^5^.

The K27M histone mutations present a therapeutic opportunity for the use of epigenetic drugs, in particular those that target chromatin-modifying proteins. Multiple publications have explored this idea, using inhibitors of histone deacetylases (HDACs)^6^, demethylases (JMJD3/UTX)^7^, methyltransferases (EZH2)^8^, and chromatin readers (BET)^9^ to demonstrate tumor regression in pre-clinical models. Clinically-translatable compounds exist to target all of these and indeed an ongoing clinical trial is testing the HDAC inhibitor panobinostat as a monotherapy (NCT02717455)^10^. However, other chromatin modifiers have yet to be explored as therapeutic targets, and there is limited investigation into how the gene expression changes generated by these drugs can be used to augment pre-existing therapies.

The histone demethylase LSD1 removes mono- and di-methyl marks from H3K4 and H3K9 and shares structural homology with monoamine oxidases (MAOs) in the brain. It can be targeted by several drug candidates^11^ and has been therapeutically exploited in other cancers such as acute myeloid leukemia^12^, sarcoma^13^, and neuroblastoma^14^. LSD1 inhibition has been shown to have an enticing therapeutic window that is selective for cancer cells, in part through its disruption of oncogenic and onco-maintenance transcriptional programs^15,16^. Furthermore, the H3K4me1 histone mark regulated by LSD1 was seen to be enriched in intergenic regions of pHGG cells^9^, suggesting that LSD1 may control access to enhancers of genes important in pHGG pathology. LSD1 inhibitors can functionally target either the catalytic domain that mediates demethylation^17^, or the scaffolding tower domain that interfaces with other proteins in epigenetic complexes^18^, and it is currently unknown what phenotype these disparate inhibitors would produce in pHGG. Given the highly disrupted yet therapeutically sensitive epigenome of pHGGs, we sought to explore in this study whether LSD1 inhibition could be both cytotoxic to pHGG and generate transcriptional changes that would inform combination therapies.

Our group previously published a report on use of a combination therapy of LSD1 and HDAC inhibition to synergistically induce cell death in adult glioblastoma cell lines and patient-derived glial stem cells^19^. In a follow-up study, we used RNA-Seq to explore how the HDAC/LSD1 inhibitor combination therapy produced gene changes in the p53 family members p63 and p73^20^. In our current study, we identify an LSD1-controlled immunogenic gene signature conserved in pHGG patients^21^ that predicts longer survival. We further show that LSD1 inhibition is selectively cytotoxic to DIPG cells, and inhibitor-based induction of this gene signature augments innate immune reactivity against DIPG by boosting natural killer (NK) cell immunotherapy response.

## Materials and Methods

Detailed experimental procedures, including CETSA, NK/T-cell cytotoxicity, drug screening and apoptosis, RT-qPCR, flow cytometry, and Western blotting, are provided in the Supplementary material.

### Cells used and assay conditions

Human pHGG cells (DIPG IV, VI, and XIII) were grown in tissue culture-treated T75 flasks (BioBasic) in TSM Base medium, defined as 50/50 DMEM/F12 medium (Corning) and Neurobasal-A medium (Invitrogen) with 1% NEAA/HEPES/sodium pyruvate/L-glutamine (Invitrogen). Before passaging or plating of cells, the following growth factors were added to TSM Base by volume: 2% B27 (Invitrogen), 0.1% of 0.2% heparin (StemCell Technologies), 20ng/mL EGF/bFGF and 10ng/mL PDGF-AA/PDGF-BB (all from Shenandoah Biotechnology). DIPG IV possess a H3.1-K27M mutation, DIPG VI and DIPG XIII are H3.3-K27M mutated. DIPG IV/VI cells were cultured as loosely-adhered monolayers (IV) or colony-forming (VI) cultures and DIPG XIII cells were cultured as free-floating neurospheres. Immortalized normal human astrocyte (NHA) cells were grown as adherent monolayer in tissue culture-treated T75 flasks in DMEM/F12 medium supplemented with 10% fetal bovine serum (Corning) and 1% L-glutamine. Human NK cells were cultured as previously described^22^. Human T-cells were isolated from buffy coats using Human T-cell Isolation Kit and cultured in ImmunoCult-XF T-cell Expansion Medium.

### Clinical datasets and bioinformatics

Gene expression in pHGGs was analyzed from the dataset published by Mackay et al.^21^ using the cBioPortal interface. The dataset was queried for a 13-gene signature identified from LSD1 knockdown RNA-Seq, and patients were clustered based on expression of 8 of the 13 genes (5 of the genes were not present in the dataset). Based on the heatmap of these 8 genes, patients were divided into low expressing (n = 142) and high expressing (n = 105). Data for fraction of the genome altered, patient age, histone mutation, anatomic location of the tumor, survival, and LSD1 expression were exported to Excel and GraphPad Prism for further analysis. CIBERSORT analysis was performed as previously described^23^ using the standard LM22 leukocyte signature matrix in the same patient dataset^21^.

### Statistical analysis

Patient data was analyzed using Wilcoxon and Log-Rank tests for survival. RT-qPCR data was analyzed using ANOVA correcting for multiple comparisons by use of the False Discovery Rate (FDR) approach. Discovery was determined using the Two-stage linear step-up procedure of Benjamini, Krieger and Yekutieli, with Q = 1%. NK and T-cell cytotoxicity was analyzed using T-tests correcting for multiple comparisons using the same FDR approach and cutoff. Comparisons were made to DMSO controls where appropriate or among each data set.

## Results

We previously performed RNA-Seq^20^ on LN18 adult glioblastoma cells when LSD1 was knocked down with shRNA nucleofection in order to explore the mechanism of their sensitivity to dual LSD1 and HDAC inhibition. In the LSD1 shRNA group alone, we applied a 1.5-fold change filter and analyzed the remaining genes with DAVID pathway analysis (Fig 1A). The 3rd-most significantly changed pathway was “immune response”, with 24 genes upregulated and downregulated by LSD1 knockdown compared to a scramble control. We sought to validate these gene changes in LN18 cells, and replicated LSD1 knockdown in the cells using electroporation and confirmed knockdown with western blot. Expression of the 13 most upregulated genes was measured with RT-qPCR and we observed a significant increase in the gene expression signature with LSD1 knockdown (Fig 1B). This confirmed our RNA-Seq data that LSD1 controls expression of these genes in a GBM cell line. Furthermore, this gene signature matches treatment of LN18 with the established LSD1 inhibitor tranylcypromine (TCP) (Fig 1B), and TCP treatment of LN18 compared with DIPG cells was non-significant (Fig 1C) indicating concordance of these upregulated immune genes between pediatric and adult glioma models. To determine the significance of this signature to patient treatment, we next proceeded to probe a dataset of 247 pediatric high-grade glioma patients (Fig 1D). Expression of LSD1 was significantly lower in patients with high expression of our gene signature, suggesting that LSD1 can control expression of these genes in pHGG patients (Fig 1E). We found our gene signature of immune response genes could predict significantly improved 5-year survival in all tumors (Fig 1F). The overall benefit was driven by K27M midline (n = 23) and WT hemispheric (n = 57) tumors; notably, this survival benefit did not extend to K27M brainstem (n = 49) tumors, and we lacked statistical power in WT brainstem (n = 9) and WT midline (n = 14) tumor samples to make strong conclusions (Fig 1F).

**Fig 1.**
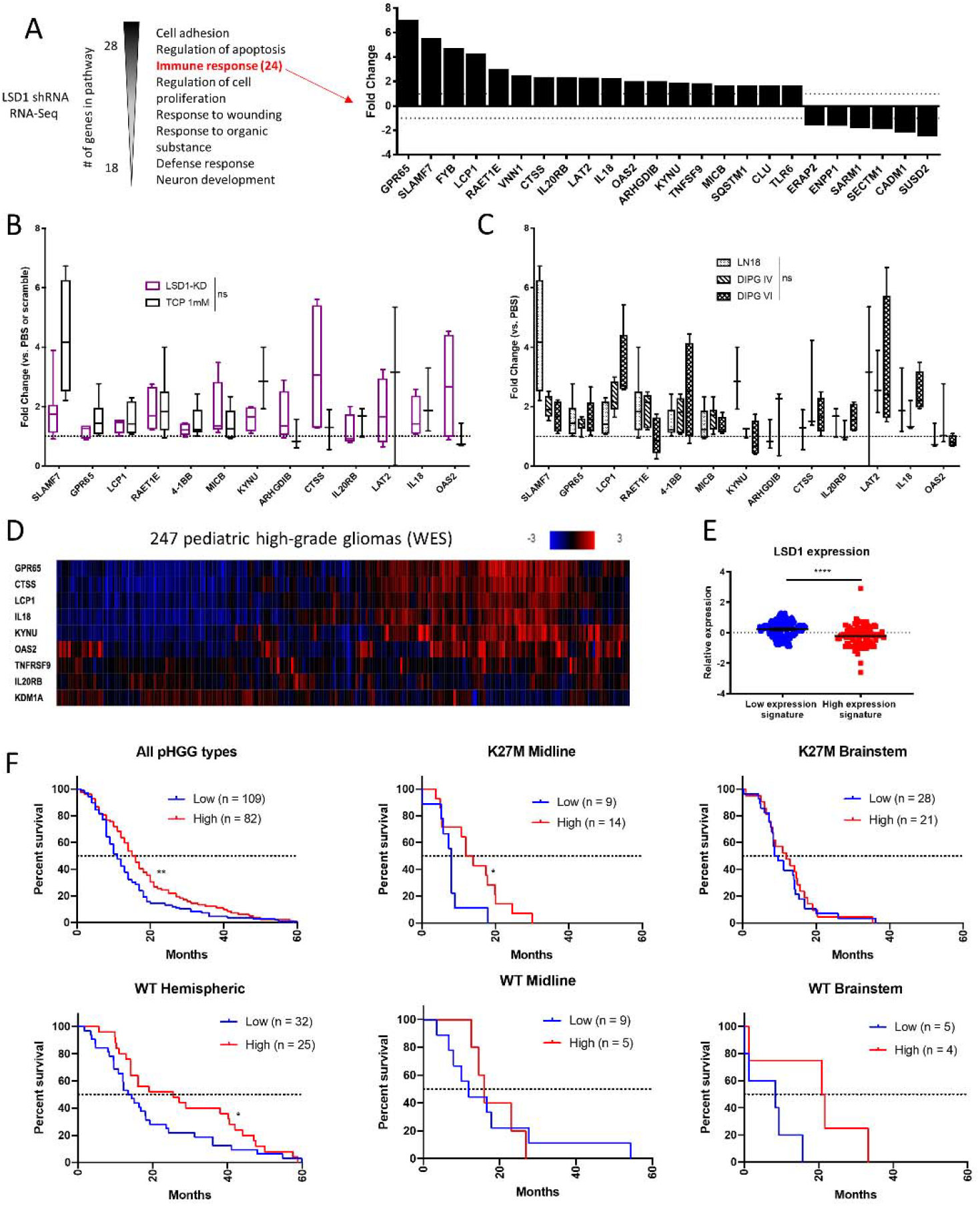
LSD1 immunogenic signature is predictive of survival benefit in pediatric high-grade glioma patients. (A) RNA-Seq pathway analysis performed in LSD1 shRNA transduced LN18 cells. Immune response genes and associated fold changes are shown. (B) RT-qPCR of immune gene signature in LN18 cells with LSD1 shRNA or 1mM TCP treatment for 24h analyzed by one-way ANOVA with FDR correction. (C) RT-qPCR of immune gene signature in LN18, DIPG IV, and DIPG VI after 1mM TCP treatment for 24h analyzed by one-way ANOVA with FDR correction. (D) Heat map of pHGG patient exome data probed for LSD1 immune gene signature. (E) LSD1 expression of patients expressing high and low levels of gene signature analyzed by unpaired T-test. (F) Survival curves of pHGG patient data subdivided by histone mutation and tumor location and analyzed by Log-Rank or Wilcoxon tests. * = p < 0.05, ** = p < 0.01, **** = p < 0.0001, ns = not significant. At least 3 biological replicates were used for RT-qPCR experiments.

In order to explore the potential of therapeutically triggering this gene signature, we profiled the potency of 3 irreversible catalytic LSD1 inhibitors (tranylcypromine, also known as TCP, GSK LSD1, and RN-1) and 2 reversible scaffolding LSD1 inhibitors (SP-2509 and SP-2577). As we have previously published, LSD1 inhibition alone in adult GBM cells does not potently reduce viability^19,20^. In pHGG cells, the same inhibitors display much greater potency that correlates with their specificity and sensitivity for inhibition of LSD1 over the related proteins LSD2, MAO-A, and MAO-B. We observed highly similar IC50s between the unique DIPG cell types for each LSD1 inhibitor tested (Fig 2 A-C). While AlamarBlue screening is a sensitive assay for cell proliferation, it cannot determine if drugs are cytostatic or cytotoxic. Therefore, we used trypan blue and DNA fragmentation assays to quantify cell death at the IC50s observed with AlamarBlue (TCP: ~1.5mM, GSK LSD1: ~400µM, RN-1: ~60µM, SP-2509/2577: ~13µM). Cell death was selectively induced in DIPG cells over normal human astrocytes (NHA) beginning at 3 days post-treatment (Fig 2D). For neurosphere forming DIPG XIII cells, we adapted another high-throughput technique to quantify cell death by use of the DNA-binding dye GelGreen and observed the same effects. In order to compare the effect of the K27M mutation on LSD1 inhibitor sensitivity, we used a murine model system where the tumor cells possess either H3.3-WT (PHC-HA) or H3.3-K27M (PKC-HA) on a shared *TP53*-flox/*PDGFRA*-overexpression background initiated in Nestin(+) neural stem cells. Both PKC-HA and PHC-HA display similar sensitivity to both LSD1 inhibitor classes, so we concluded the growth inhibitory effects of LSD1 inhibition to be K27M-independent (Fig 2E).

**Fig 2.**
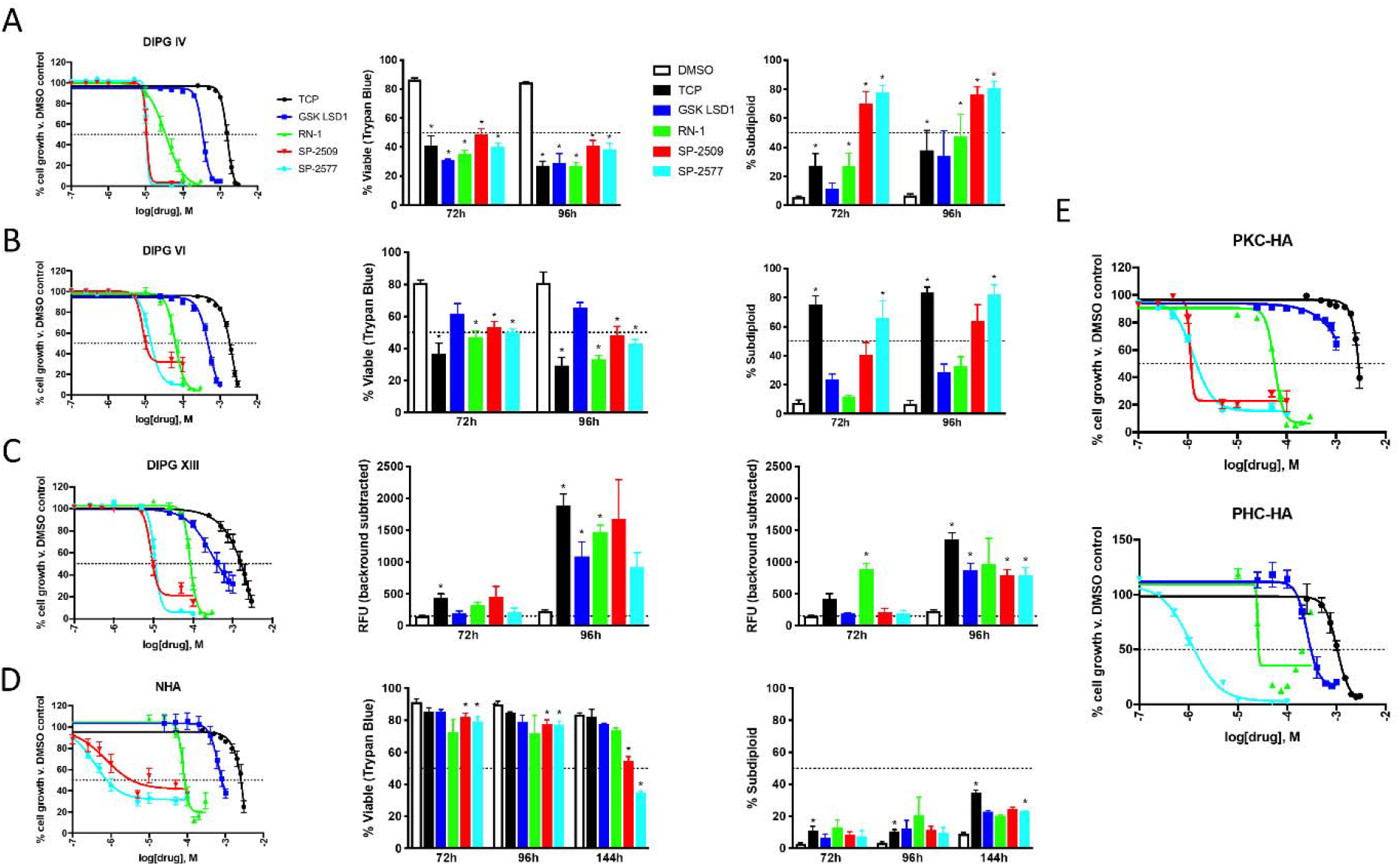
LSD1 inhibitors are growth inhibitory and induce selective cell death in DIPG cells. (A) Dose response curves of LSD1 inhibitors in DIPG IV, (B) DIPG VI, (C) DIPG XIII, (D) NHA, and (E) PKC/PHC-HA cells measured using AlamarBlue after 120h treatment. Cell viability after 72 and 96h measured using trypan blue cell exclusion and analyzed by T-test comparing with DMSO control using FDR correction. DNA fragmentation measured using propidium iodide on flow cytometry analyzed by T-test comparing with DMSO control using FDR correction. Cell death of DIPG XIII (C) measured using GelGreen fluorescent intensity in 96-well plate reader and analyzed by T-test comparing with DMSO control using FDR correction. * = p < 0.05. At least 3 biological replicates were used for all experiments. Error bars represent mean +/− SEM.

We further profiled the on-target binding of our LSD1 inhibitor suite through use of the cellular thermal shift assay (CETSA). By heating live cells under treatment with candidate compounds and interrogating the thermostability of the target protein via western blot (Fig 3A), we could determine if LSD1 is binding to our LSD1 inhibitors in DIPG and NHA cells (Fig 3B). It was observed that all catalytic LSD1 inhibitors could bind LSD1 in all cell types, while results were less consistent with the scaffolding LSD1 inhibitor compounds (Fig 3C). We hypothesized our dose of SP-2509 and SP-2577 may be too low to thermostabilize LSD1, so we conducted dose response CETSAs with TCP as a positive control. We found some increase in binding by increased doses of SP-2509, but no increase in binding with higher doses of SP-2577 (Fig 3D). Given that we dosed up to 100 µM for the dose response CETSA, which is almost 10X the IC50 of the scaffolding inhibitors in DIPG cells, either the CETSA assay cannot capture the protein complex-disruption properties of the scaffolding compounds or there exists off-target effects, of which there is published data for rationale of the latter^25,26^.

**Fig 3.**
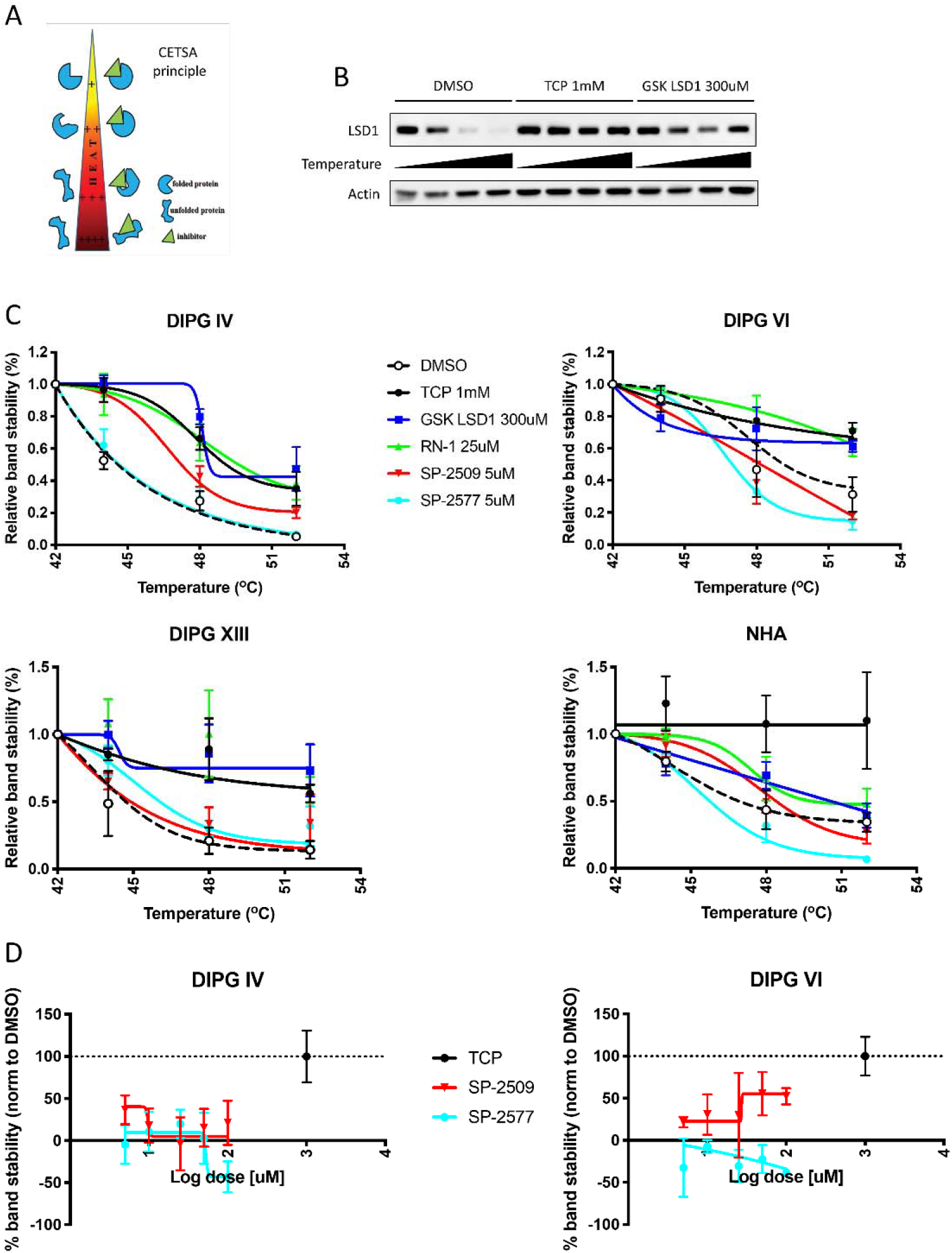
Irreversible catalytic LSD1 inhibitors thermostabilize LSD1 in DIPG and NHA cells. (A) Principle of the cellular thermal shift assay (CETSA). Figure adapted from Hau et al., Epigenetics, 2017. (B) Representative western blot of CETSA probing LSD1 thermostability in DIPG VI cells. (C) Protein melt curves for LSD1 in different cell types. Each data point was normalized to beta-actin level and further normalized to 42C data point for each experimental condition. (D) Dose response CETSA for scaffolding inhibitors SP-2509/2577. LSD1 was destabilized at 48C for all doses and DMSO control was set as 0% stability and 1mM TCP was set as 100% stability. At least 3 biological replicates were used for all experiments. Error bars represent mean +/− SEM.

With sensitivity and on-target activity of LSD1 inhibition in DIPG established, we next treated cells with sub-cytotoxic doses of LSD1 inhibitors for 24h and isolated RNA to measure expression of our immune gene signature. DIPG cells display a significant upregulation of the signature under treatment with irreversible catalytic LSD1 inhibitors, but no significant changes when treated with reversible scaffolding LSD1 inhibitors SP-2509 and its clinical successor SP-2577 (Seclidemstat). This gene signature was also selective for DIPG, as the same treatment did not induce upregulation NHA cells (Fig 4A-B). Several genes in the signature correspond to immune signaling receptors, so we next profiled protein expression of 3 innate immune receptors (SLAMF7, MICB, and ULBP-4) using flow cytometry. DIPG cells display differing baseline levels of these receptors, but overall, we could detect increased expression on live cells after LSD1 inhibitor treatment for 48h (Fig 4C).

**Fig 4.**
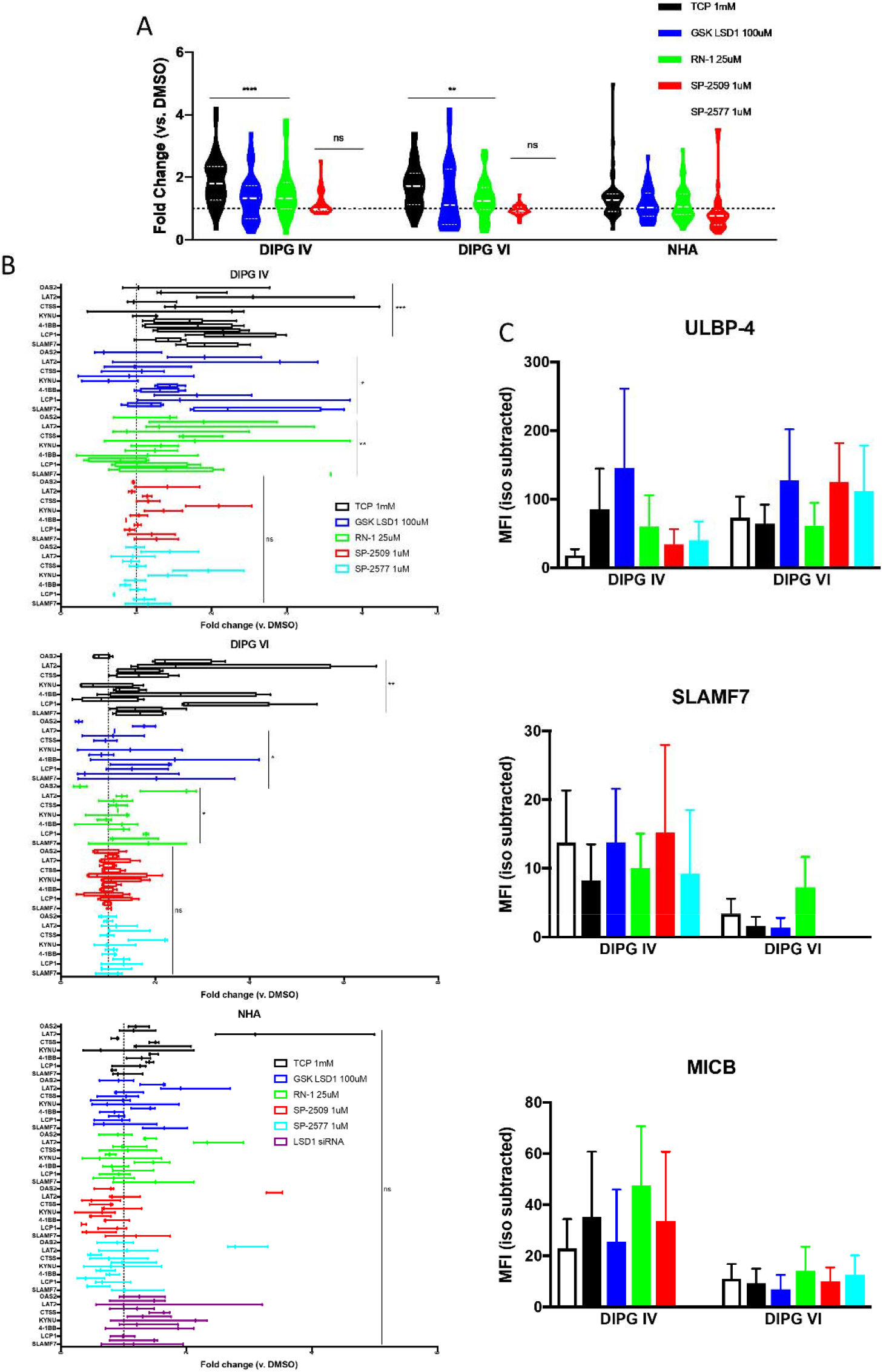
Irreversible catalytic LSD1 inhibitors selectively generate immunogenic signature in DIPG cells. (A) RT-qPCR for immune gene signature performed on cells after treatment with indicated LSD1 inhibitors for 24h. Catalytic inhibitors (TCP, GSK LSD1, and RN-1) and scaffolding inhibitors (SP-2509/2577) are compared to matched NHA controls using one-way ANOVA with FDR correction. (B) RT-qPCR data re-plotted with individual genes and including siRNA treatment for 48h. Fold change compared to DMSO control analyzed via one-way ANOVA with FDR correction. (C) Median fluorescent intensity of indicated receptors after 48h of LSD1 inhibitor treatment. Matched species and fluorophore isotype controls used to measure background fluorescence. * = p < 0.05, ** = p < 0.01, *** = p < 0.001, **** = p < 0.0001, ns = not significant. At least 3 biological replicates were used for all experiments. Error bars represent mean +/− SEM.

The above receptors are known to play roles in NK cell signaling, either as NKG2D ligands or self-ligating receptors, and stimulation through these receptors increases NK cell activity and lysis of target cells. Given our observed upregulation of these receptors under LSD1 inhibition, we hypothesized that NK cells would lyse target DIPG cells more readily when LSD1 is inhibited. Using fluorescently labeled DIPG cells, we incubated them with effector human NK cells at various effector to target (E:T) ratios. Across multiple unique healthy blood donors, we could observe increases in lysis in 2 DIPG lines when treated with catalytic LSD1 inhibitors TCP and GSK LSD1, but inconsistently under scaffolding LSD1 inhibition by SP-2509 (Fig 5A). Notably, the lysis efficacy of expanded healthy human donor T-cells was much lower than NK cells, and could not be augmented by LSD1 inhibitor pre-treatment (Fig 5B). We aimed to correlate genetic biomarkers of NK lysis by probing our gene signature from matched co-culture samples, and observed strong positive trends for 4 genes in DIPG IV (Fig 5C) and 2 genes in DIPG VI (Fig 5D). Unexpectedly, a negative correlation could be found for 4-1BB (Fig 5E), traditionally a T-cell stimulatory factor, which could indicate alternative function during NK cell engagement.

**Fig 5.**
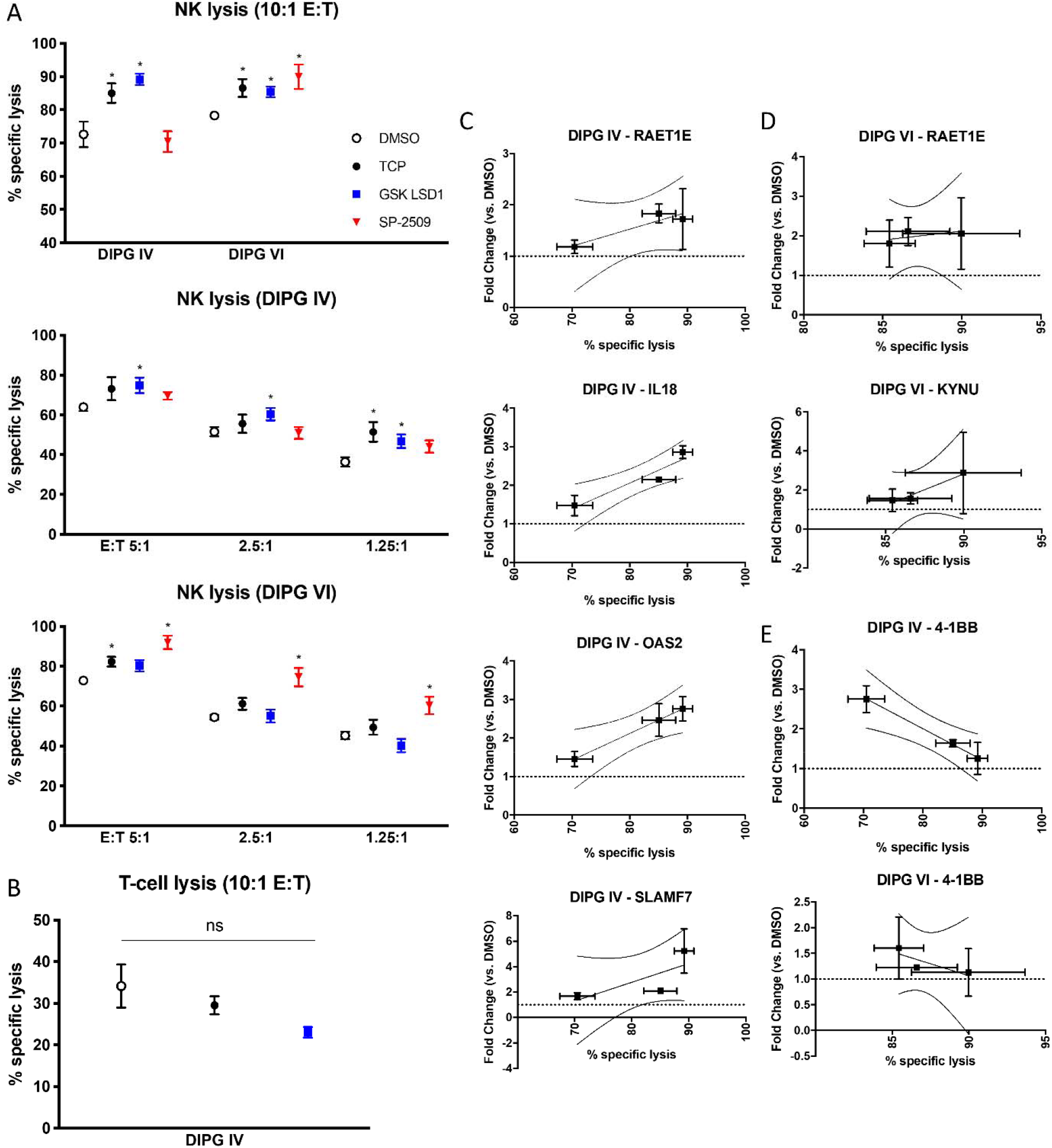
LSD1 inhibition upregulates innate immune receptors and sensitizes DIPG cells to NK cell lysis which correlates with unique genetic identifiers of response. (A) Lysis of target DIPG cells co-cultured with NK cells after 48h pre-treatment of target cells with LSD1 inhibitors (TCP 0.5mM, GSK LSD1 300µM, SP-2509 5µM). Treatments analyzed versus DMSO control using T-test with FDR correction. (B) Lysis of target DIPG cells co-cultured with T-cells after 48h LSD1 inhibitor pre-treatment. (C) DIPG IV RT-qPCR from matched co-culture experiments, genes with positive Pearson’s correlation R^2^ > 0.80 are shown with 95% confidence intervals. (D) DIPG VI RT-qPCR from matched co-culture experiments. (E) RT-qPCR from matched co-culture experiments, negative correlation with R^2^ > 0.80 shown with 95% confidence intervals. * = p < 0.05 and ns = not significant. At least 3 biological replicates were used for all experiments. Error bars represent mean +/− SEM.

To further validate our finding that catalytic LSD1 inhibition can enhance NK cell lysis of DIPG, we re-visited our patient data for analysis using CIBERSORT, an *in silico* method to determine immune cell infiltrate in bulk sequenced tissue. We found that significant NK cell infiltration is positively prognostic for H3-WT hemispheric tumors, but CD8 T-cell infiltrate is negatively prognostic. Brainstem tumors benefited less from NK infiltrate, but significant NK presence was still a superior prognostic indicator versus CD8 T-cells in the brainstem (Fig 6A). We next hypothesized how these already-present immune cells would respond to LSD1 inhibition, and treated expanded NK and T-cells with a panel of epigenetic inhibitors including our LSD1 suite. As has been known, T-cells are sensitive to HDAC inhibition, but are fairly resistant to LSD1 inhibition except at higher doses of the scaffolding inhibitors. Conversely, NK cells are resistant to HDAC inhibition but highly sensitive to scaffolding LSD1 inhibitors, with no live cells detected even at 0.5µM doses of SP-2509/2577. Catalytic LSD1 inhibitors are comparatively non-perturbing, with the IC50s against NK cells being 2-10X higher than doses needed to induce our gene signature (Fig 6B). Given our data showing the scaffolding LSD1 inhibitors are cytostatic but not cytotoxic to NHA cells, we profiled the metabolism of both NK and T-cells after LSD1 inhibitor treatment, as active metabolism of nutrients has been shown to be crucial to anti-tumor effects of both cell types. Strikingly, the scaffolding LSD1 inhibitors completely suppress the metabolism of NK cells, rendering them metabolically quiescent but still alive at 48h post-treatment (Fig 6C). Collectively, this data suggests that catalytic LSD1 inhibitors can be used at therapeutic doses to induce increased NK cell reactivity without harming the NK cells directly.

**Fig 6.**
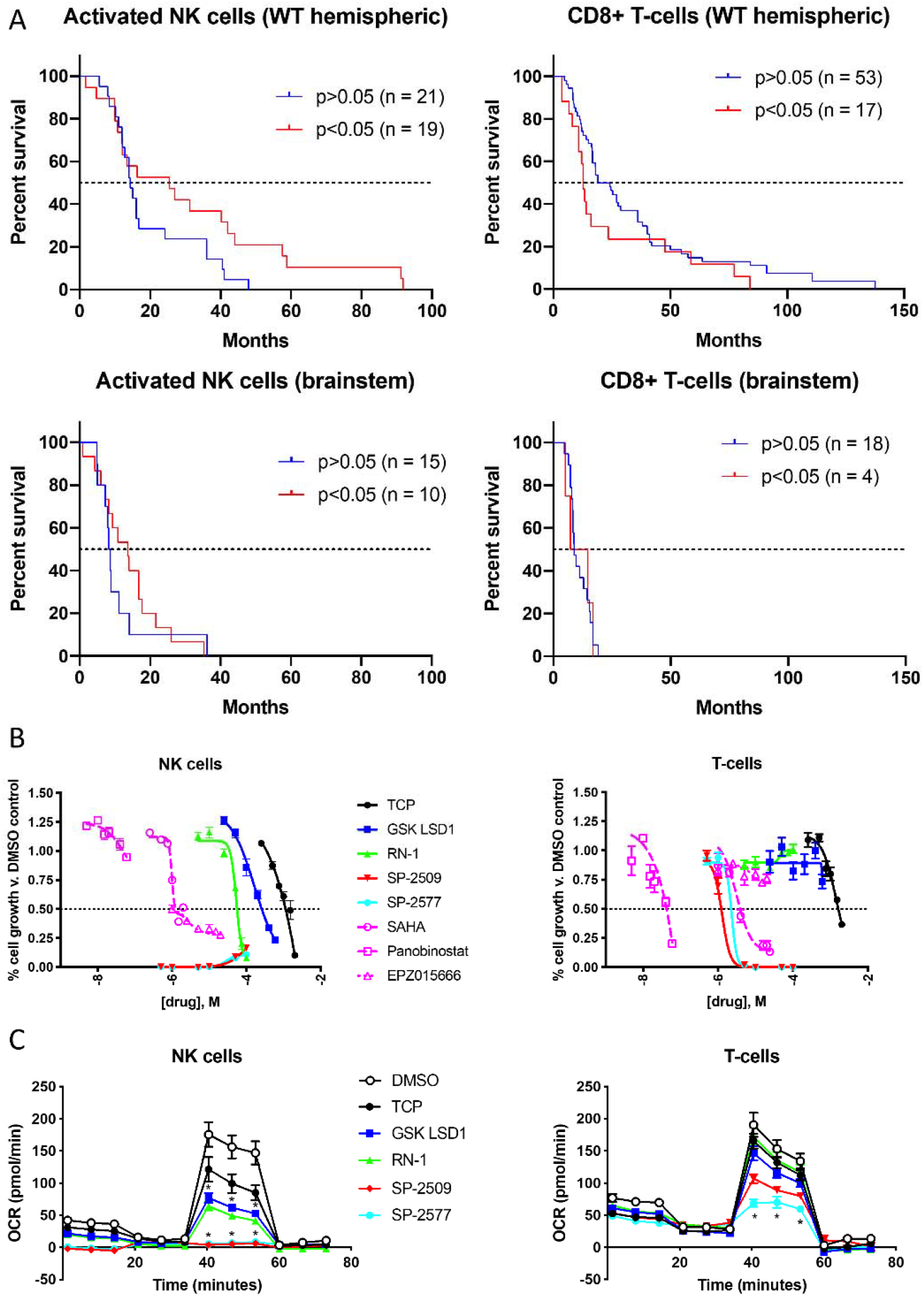
NK cell tumor infiltration is predictive of survival benefit in pediatric high-grade glioma patients and catalytic LSD1 inhibitors are non-perturbing to mature NK and T-cells. (A) CIBERSORT analysis of pHGG patient data sub-analyzed by tumor location and immune cell type. Survival curves show significant vs. non-significant presence of indicated immune cell in patient tissue. (B) Purified expanding T-and NK cells treated with indicated epigenetic inhibitors for 120h and measured using AlamarBlue. (C) XF Mito Stress Test performed on NK and T-cells after 48h of LSD1 inhibitor treatment (TCP 0.5mM, GSK LSD1 300µM, RN-1 25µM, SP-2509/2577 5µM) and treatments compared to DMSO control analyzed by T-test with FDR correction. * = p < 0.05. At least 3 biological replicates or unique donors were used for all experiments. Error bars represent mean +/− SEM.

## Discussion

Here we have described a novel dual role for the histone demethylase LSD1 in controlling cell death and innate immune responses against pHGG and DIPG cells. Expression of immune response genes accurately categorized pHGG patients’ survival probability, and therapy with catalytic LSD1 inhibitors triggered gene expression changes that enhanced NK cell-mediated lysis. Epigenetic drugs of various targets have shown promising anti-tumor effects in pre-clinical DIPG models, but these agents have not yet been combined with clinically relevant immunotherapy regimens. NK cell administration is effective against malignant gliomas^27^, can be grown without autologous sources of leukocytes^28^, and has already begun clinical trials using our exact NK-expansion methodology^29^. Furthermore, combination therapy with LSD1 inhibitors is desirable due to the demonstrated selectivity of LSD1i-induced DIPG cell death versus normal astrocytes. These results suggest the clinical potential of a regimen combining irreversible catalytic LSD1 inhibitors and NK cell infusion in pHGG.

LSD1 recently was shown to control the dsRNA stress response and subsequently synergize with anti-PD1 therapy in murine models^30,31^. Notably, this phenomenon was described only in epithelial-derived tissue such as breast, skin, and lung cancer cell lines. We could not recapitulate these genetic changes in our LSD1 shRNA RNA-Seq and believe this aspect of LSD1 influence on tumor immunity may be restricted to only certain cell types. Furthermore, these investigations did not thoroughly explore use of LSD1 inhibitors with differential mechanisms of action to induce enhanced adaptive immunity. DIPG and its microenvironment has recently been described as “immune cold” to the adaptive immune system but potentially vulnerable to modulation of innate immunity ^32–34^. Another consideration is the pharmacokinetic (PK) properties of currently available LSD1 inhibitors, which have recently been shown to be ineffective against intracranial medulloblastoma despite very potent *in vitro* activity^35^. Two separate groups in Japan recently described new catalytic LSD1 inhibitors with favorable PK properties for potential use in neuro-oncology^36,37^. The transcription factor GFI1B is not disrupted by their compounds, reducing peripheral toxicity and sparing immune cells that would be critical for our combination therapy. Furthermore, they target the LSD1 catalytic site and may be able to induce the NK-boosting gene changes seen in our experiments.

NK cells have been shown to specifically target glioblastoma stem cells^38^ and can circumvent antigen loss seen with CAR-T-based modalities^39^. A clinical trial of NK cell infusion for pediatric patients with CNS tumors is already in-progress^40^ and preliminary results indicate an excellent safety profile. Other promising clinically-actionable drug candidates in DIPG could also be combined with NK cell therapy, such as the DRD2-antagonist ONC201 which has been shown to recruit NK cells into tumors^41^, and has clinical trial data in adult gliomas^42^ and recently began a trial in pediatric K27M+ tumors (NCT03416530). Oncolytic viruses targeting DIPG have also begun clinical trials^43^ and NK cells have been shown to be involved in the immune response to lysed tumor cells in the brain^44^. We did not yet test synergy of NK cells with standard-of-care radiotherapy or chemotherapy in our models, which is warranted based on pre-clinical^45,46^ and clinical trial^47^ data of adult gliomas.

Targeted immunotherapy of DIPG was recently demonstrated through use of CAR T-cells directed against the cell surface glycolipid GD2^48^. Impressive tumor regression was achieved; however, toxicity was observed driven by excessive cranial inflammation in the brainstem and thalamus. A similar phenomenon was observed in GD2-directed CAR-T in neuroblastoma^49^, suggesting that highly-potent immunotherapies may be inadvisable in infratentorial tumors and combination therapies that boost native immune responses may be a safer alternative. While pediatric gliomas possess an overall low mutational burden, it was previously unknown if patients could generate acquired immunity against neoantigens specific to pHGG and DIPG. Chheda et al.^50^ demonstrated that DIPG triggers a T-cell response directed against the H3.3-K27M epitope, and this T-cell receptor can be cloned into autologous T-cells for antigen-specific immunotherapy. A K27M-transfected U87 model was used to show in vivo efficacy with no encephalitis^50^, but it remains unclear if the lack of encephalitis was due to the U87 cell type or if H3.3-K27M-TCR-transduced T-cells do not induce the uncontrollable inflammatory response seen with CAR-T therapy.

The novel findings in our paper set the stage for further investigations into innate immune interactions in DIPG and how they can be therapeutically exploited with epigenetic inhibitors that can induce both cell death and NK cell reactivity. Our extensive use of LSD1 inhibitors with unique binding properties suggests future investigation of combining brain-penetrant catalytic LSD1 inhibitors and NK cell infusions for synergistic killing of DIPG tumor tissue.

## Acknowledgements

The authors thank Drs. Michael Curran, Candelaria Gomez-Manzano, Andrew Gladden, and Min Gyu Lee for their input on experimental design and directions of the study. They thank Drs. Ryan G. Kruger and Helai P. Mohammed for experimental suggestions and additional supply of GSK LSD1. They thank Drs. Ruolan Han and Jeffrey L. Larson for providing SP-2577 and data interpretation. They also wish to thank Drs. Vidya Gopalakrishnan and Aundrietta Duncan for helpful discussions and data interpretation.

