## Supplementary Material for "Pharmacologic inhibition of lysine specific demethylase-1 (LSD1) as a therapeutic and immune-sensitization strategy in diffuse intrinsic pontine glioma (DIPG)"

**Supplementary Methods**

*Cells and human samples*

DIPG and NHA cells were validated at least once per year by STR DNA fingerprinting using the Promega 16 High Sensitivity STR Kit (Catalog # DC2100). The STR profiles were compared to online search databases (DSMZ/ATCC/JCRB/RIKEN) of approximately 2500 known profiles; along with the MD Anderson Characterized Cell Line Core (CCLC) database of approximately 2600 know profiles.

All adherent or neurosphere cells were detached and/or dissociated with TrypLE during normal passage or Accutase during analysis or use in experiments. All cell lines were cultured without antibiotic and monitored for mycoplasma with MycoAlert PLUS (Lonza) with luminescence being read on a Synergy 2 plate reader (BioTek). Cell lines were cultured for an average of 3 months after being thawed, with mycoplasma testing done after thawing and prior to freezing to maintain myco-free stock, as well as periodically during experimental periods.

NHA cells were additionally cultured with 0.5 µg/mL puromycin (Sigma) and 10 µg/mL blasticidin (Cayman Chemical) to maintain a transformed E6/E7/TERT-overexpressing phenotype. NK cells were isolated from buffy coats using RosetteSep Human NK cell Enrichment Cocktail (StemCell Technologies). After isolation, cells were stimulated with membrane-bound IL-21–expressing artificial antigen-presenting (mbIL-21 aAPC) K562 expansion cells (irradiated at 100Gy) at a ratio of 2:1 aAPC:NK in RPMI 1640 medium (Corning) supplemented with 10% FBS and 1% L-glutamine/NEAA/HEPES/sodium pyruvate and additionally supplemented with 50 U/mL IL-2 (StemCell Technologies) replenished every 3 days. T-cells were supplemented with 50 U/mL IL-2 and stimulated for expansion with Human CD3/CD28/CD2 T-cell activator (StemCell Technologies) every 7 days.

*Cellular thermal shift assay (CETSA)*

At least 1 x 10^6^ NHA or DIPG cells were plated in T75 flasks for each experimental condition and treated with LSD1 inhibitors for 1 hour. Cells were then harvested with TrypLE, washed in PBS, and resuspended in 200 μL cold CETSA wash buffer (defined as PBS with protease inhibitor cocktail added). 50 μL of each experimental condition were aliquoted into PCR strip tubes to make the melt curve. For LSD1, the temperatures were 42, 44, 48, and 52 C; this will vary for each protein being interrogated. Each set of aliquots was heated in a gradient thermocycler (BIO-RAD) for 3 mins then cooled to 25C indefinitely. Strip tubes were immediately freeze/thawed in liquid nitrogen for 3 cycles to induce cell lysis. Lysates were spun down in a microcentrifuge at 12,000 RPM for 20 mins at 4C. Cleared lysates were either frozen at -80C or 15 μL was immediately loaded onto polyacrylamide gels for Western blot as described.

*NK and T-cell cytotoxicity co-culture*

Target cells were grown under treatment conditions for defined times and doses, then harvested with Accutase and stained with calcein AM at 4 µM in NK cell media for 60 mins at 37C. Calcein AM-stained cells were counted and plated in 96-well round bottom plates (Corning) at 50,000 cells/well, then NK or T-cells were added at defined effector-to-target ratios. 1% Triton-X (max lysis) and media only (background lysis) of target cells alone were included for each treatment condition. Plates were spun down at 100 x g for 1 min to initiate cell contact and then incubated for 4 hours at 37C. Following incubation, wells were mixed gently and plates then spun down at 100 x g for 5 mins, and 100 µL supernatant media was moved to clear-bottom, white-walled 96-well plates. Fluorescence was read at 485nm excitation/530nm emission on a Spectramax Gemini EM plate reader (Molecular Devices) with bottom read setting.

*Chemicals and antibodies*

The compounds tranylcypromine (TCP) (Enzo Biosciences), GSK LSD1 (Cayman Chemical), RN-1 (Cayman Chemical), and SP-2509 (EMD Millipore) were purchased from the indicated vendors. SP-2577 was provided as a free base formulation by Salarius Pharmaceuticals. Additional amounts of GSK LSD1 were supplied by GlaxoSmithKline. TCP was suspended in phosphate-buffered saline solution (PBS), while all other drugs were suspended in dimethyl sulfoxide (DMSO) and aliquoted for storage at -20^o^C. AlamarBlue was made from 2 g resazurin sodium salt (Sigma) resuspended in 500mL sterile PBS and stored at 4^o^C as a 100x solution. GelGreen (Biotium) was stored in the dark at room temperature. Calcein AM (EMD Millipore) was resuspended in DMSO and stored at -20^o^C. Ghost Dye Red 780 and Ghost Dye Violet (Tonbo Biosciences) were aliquoted for storage at -20^o^C.

Antibodies for LSD1 (Abcam), β-Actin (Sigma), SLAMF7 PE (Biolegend), MICB APC (R&D Systems), and ULBP-4 Alexa 488 (R&D Systems) were used at manufacturer recommended dilutions for western blot or flow cytometry. Isotype antibodies matched to the species, class, and fluorophore were used in flow cytometry experiments. UltraComp eBeads (ThermoFisher Scientific) were used to calculate compensation for multi-color flow cytometry experiments.

*Drug screening*

LSD1 inhibitors were screened for efficacy against cells using 96-well flat-bottom plates (BioBasic) and AlamarBlue fluorescence as readout for live cell number or GelGreen fluorescence as readout for cell death^24^. Cells were plated as single cells at 5,000 cells/well in 150µL of medium and were allowed to adhere overnight (NHA/DIPG IV) or were grown for 3-4 days until colonies (DIPG VI) or neurospheres (DIPG XIII) formed. Only the inner 60 wells of the plate were used; wells on the perimeter were filled with 200 µL PBS to control for edge effect. For treatment, drugs were diluted in medium to a 6X working stock, and 30 µL of the stock was added to the 150 µL of medium in each well for a total of 180 µL/well. For live cell counts, plates were incubated for 4 days, and 18 µL AlamarBlue was added at the end of day 4. On day 5, fluorescence was read at 540nm excitation/600nm emission on a Synergy 2 plate reader with the bottom read setting. For cell death count, cells were plated in white-walled flat clear-bottomed 96-well plates (Grenier) and grown as above. GelGreen was added during drug treatment to a final concentration of 2X, and fluorescence was read at 485nm excitation/528nm emission as above. Using GraphPad Prism 8.1.2, dose responses were transformed to log scale and normalized to DMSO controls; a sigmoidal curve was plotted to calculate the median inhibitory concentration (IC_50_).

*Cell viability and apoptosis assays*

For confirmation of cell death following AlamarBlue screening, cells were plated and treated at the IC50 doses determined from their respective sigmoidal curves. After 72 or 96 hours, cells were harvested with TrypLE, spun down, and resuspended in 800 µL PBS. 500 µL of cells were analyzed for viability by TrypanBlue exclusion on a ViCell XR (BeckmanCoulter). The remaining 300 µL was fixed by adding 700 µL dropwise of ice cold 100% ethanol and storing at -20C. After a minimum of 24 hours, cells were spun down, washed in PBS, and resuspended in a mixture of 300 µL PBS with 37.5 µM propidium iodide and 100 µg/mL of Ribonuclease A and incubated for 30 mins at RT in the dark. Cells were immediately analyzed on a Fortessa flow cytometer and subdiploid populations were quantified in FlowJo.

*RNA isolation and RT-qPCR*

RNA was isolated using the RNeasy kit (QIAGEN) following manufacturer protocol. RNA was quantified on a Nanodrop 1000 spectrophotometer (ThermoFisher) and 500-1000 µg of RNA was reverse transcribed into 20 µL cDNA using the iScript cDNA synthesis kit (BIO-RAD). cDNA was diluted in template buffer (Biotium) by 2X, and 1 µL cDNA was plated in duplicate or triplicate on a 96-well qPCR plate (USA Scientific) mixed with 10 µL 2X Forget-Me-Not EvaGreen qPCR Master Mix, 8 µL nuclease-free water, and 1 µL of a 10 µM mix of forward and reverse primers for genes of interest. Primers are listed in supplementary table 1. Assay was run on a LightCycler 96 instrument (Roche) using Biotium protocol and analyzed with LC96 software (Roche) to confirm amplification and single melt peaks. Ct values were exported and analyzed in Excel using the 2^-ΔΔCT^ method compared to DMSO controls. Fold changes were plotted in GraphPad Prism using multiple biological replicates.

*Flow cytometry*

Cells were harvested with Accutase after being treated for indicated time points and doses and washed with PBS in 5mL FACS tubes. Ghost Dye Red 780 was diluted 10X in PBS and this mixture was used to resuspend cells in 50 µL/tube for isotype and on-target staining. After 10 mins incubation at room temperature, 50 µL of antibody mixture diluted in 0.5% BSA in PBS was added using the manufacturer recommended dilutions of 5 µL/1 x 10^6^ cells. Tubes were incubated at 4C for 25 mins, washed with PBS, and resuspended in 300 µL 0.5% BSA in PBS for acquisition on a Fortessa flow cytometer (BD) with 488nm and 640nm laser configuration. Data was analyzed with FlowJo 10.6.0 (FlowJo, LLC) gating on live cells and measuring MFI values of indicated fluorophores versus DMSO control.

*Cell transfections*

NHA cells were transfected with a scramble (control) or LSD1 siRNA cocktail (Santa Cruz Biotechnology) using Lipofectamine RNAiMAX (ThermoFisher Scientific) with the standard protocol for 6-well plates. Cells were incubated for 48 hours then harvested for lysates and RNA. Knockdown was confirmed via western blot.

*Western blotting*

At least 1 x 10^6^ cells were harvested with TrypLE and washed once with PBS, followed by lysis with Triton-X buffer for at least 1 hour rotating at 4^o^C. Lysates were spun at 12,000 RPM for 20 minutes at 4^o^C to pellet debris. Protein content was measured via Bradford assay (BIO-RAD) with bovine serum albumin (BSA) diluted in PBS used to establish the standard curve. Absorbance was measured at 750 nm on a SpectraMax Plus 384 plate reader (Molecular Devices). Equal amounts of protein were loaded on a polyacrylamide gel for sodium dodecyl sulfate-gel electrophoresis and run at 100V for 2 hours. Proteins were transferred to polyvinylidene fluoride (PVDF) membranes via wet transfer at 100V for 1 hour. Membranes were blocked with 1% fish gelatin for 1 hour at room temperature. Antibodies were incubated overnight at 4^o^C with gentle agitation. The next day, the membranes were washed with Tris-buffered saline solution containing Tween (TBST) and incubated with horseradish peroxidase–conjugated (HRP) secondary antibody (Cell Signaling Technology). Proteins were visualized by SignalFire ECL Reagent (Cell Signaling Technology) for 1 minute and imaged on a ChemiDoc Touch (BIO-RAD). Images were evaluated with Image Lab software (BIO-RAD) and protein expression quantified with ImageJ (US National Institutes of Health).

**Supplementary Figure Legends**

Supplementary Figure S1. *Confirmation of NHA LSD1 siRNA knockdown*. NHA cells transfected with LSD1 siRNA for 48h were harvested for RNA (main manuscript) and lysates for Western blot. Example blot appears on the right and quantitation of 3 biological replicates appears on the left. LSD1 bands were normalized to actin and further normalized to “no RNA” control and plotted after quantification via ImageJ.

Supplementary Figure S2. *Healthy human NK cell donor phenotype after expansion*. NK cells were expanded as described in supplementary methods and analyzed for phenotype by FACS. Cells were pelleted, stained with Ghost Dye Violet (Tonbo) in PBS, then stained with the following antibodies at manufactured recommended concentrations in FACS buffer (PBS + 2% BSA): CD3-FITC (BD), CD8-APC-Cy7 (Biolegend), CD16-PE-Cy7 (Biolegend), CD56-PE (BD), SLAMF7-PE (Biolegend), CD4-PerCP (Thermo), and NKG2D-APC (Biolegend).

**Supplementary Figure S1**

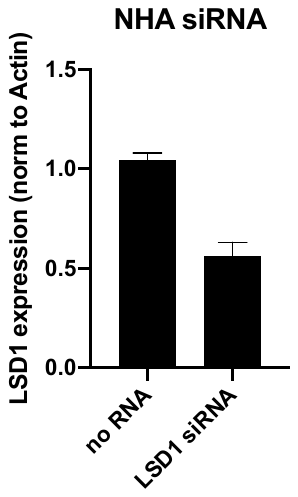

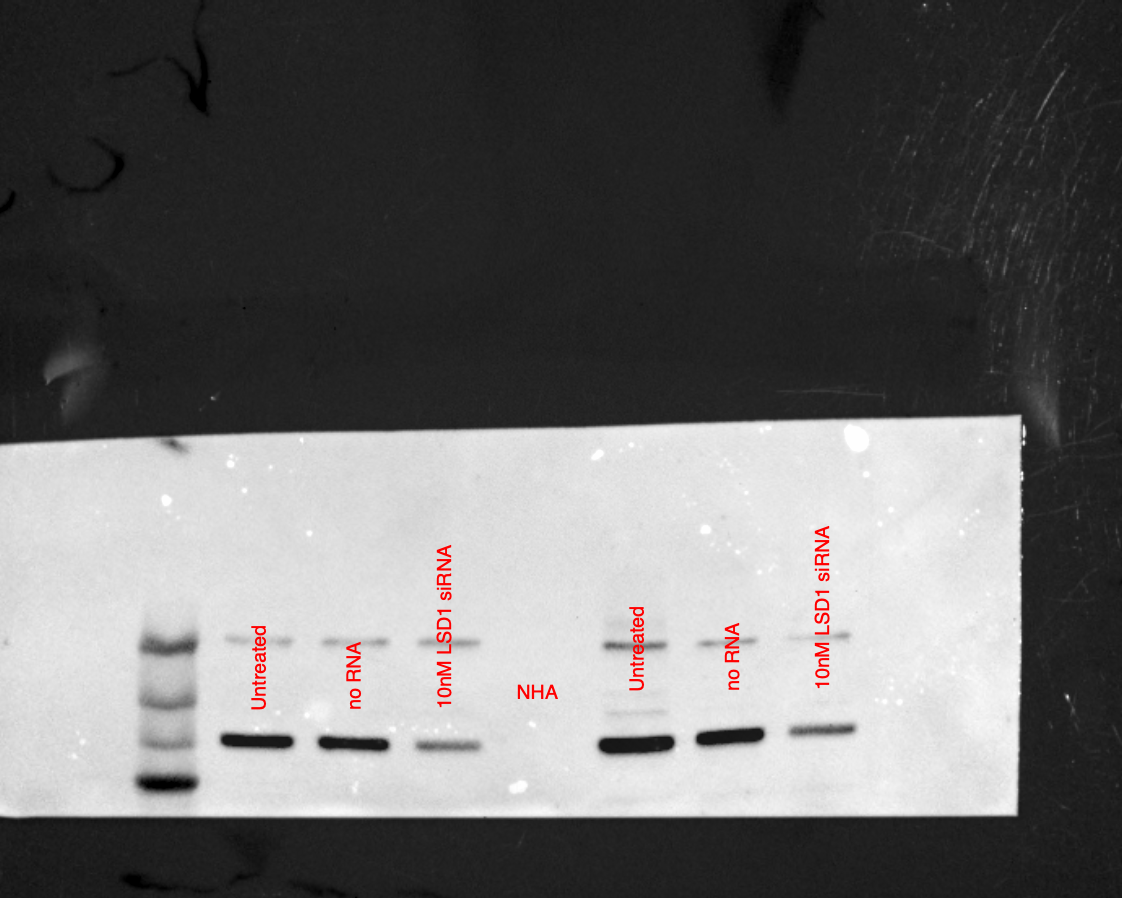

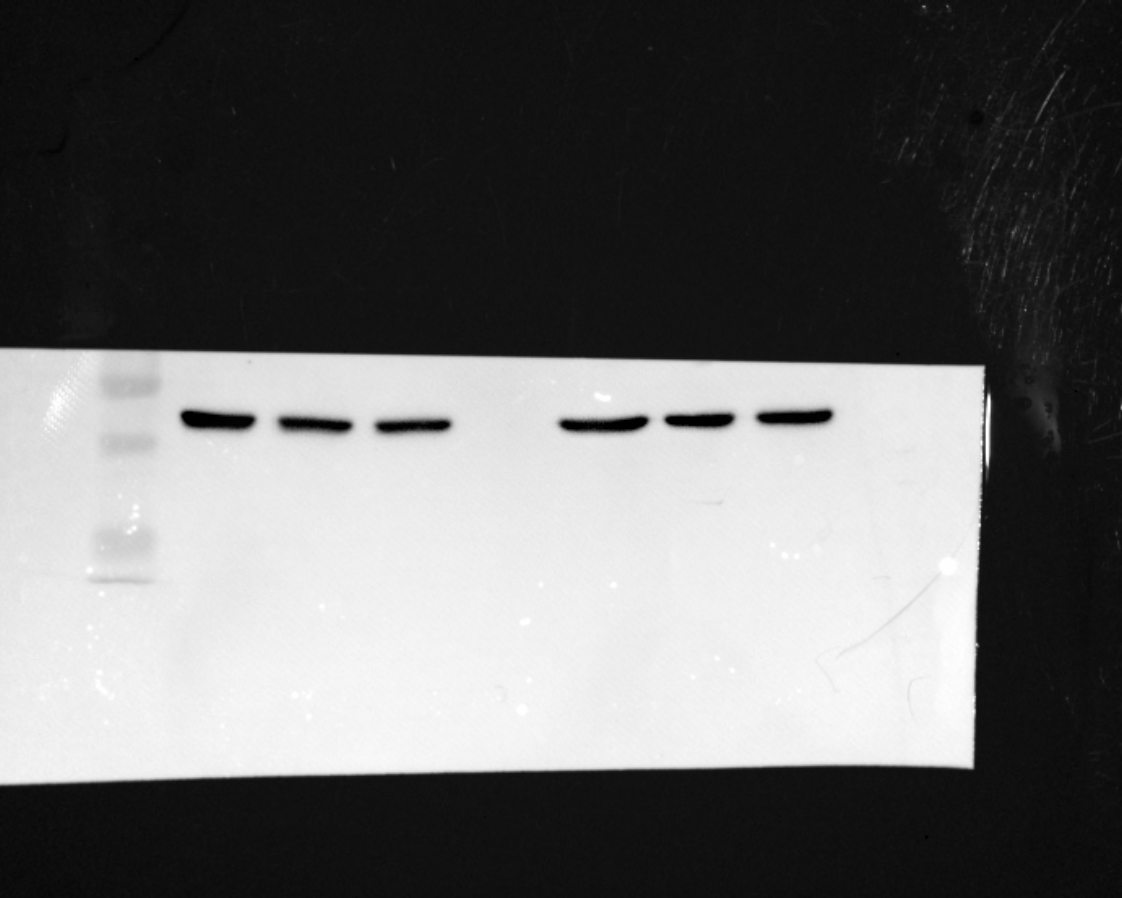

LSD1

Actin

**Supplementary Figure S2**

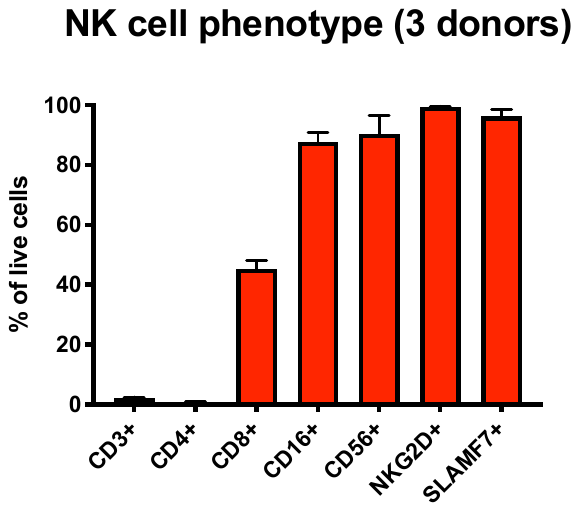

**Supplementary Table 1**

| **Primer** | **Sequence** |
| --- | --- |
| SLAMF7 Forward | AAGGGGAATGGCTGCTTTTG |
| SLAMF7 Reverse | CTCAATCCCATTCTTGCCCAAC |
| GPR65 Forward | CATCCCACCTAGGTCTCCCA |
| GPR65 Reverse | CACATCACTTCCCCCTCACC |
| LCP1 Forward | GCAGTTTGTCACAGCCACAG |
| LCP1 Reverse | TCATTGACCTTCTGGCCACC |
| RAET1E Forward | TGTGAAGCGCAGGTCTTCTT |
| RAET1E Reverse | AACAGGATGAATGCCCCCAG |
| 4-1BB Forward | TGCTTGTGAATGGGACGAAG |
| 4-1BB Reverse | ACGTCAGCGCAAGAAAGAAG |
| MICB Forward | ATGAGGTGTTTGCTGCTCTG |
| MICB Reverse | TTTGCCCACATCCTGCATTC |
| KYNU Forward | TCAGTGGAGACCATCGACAG |
| KYNU Reverse | GCATTTGAGTTCAGCCGCAA |
| ARHGDIB Forward | GCCCAGGGTTTCCTCTTCAA |
| ARHGDIB Reverse | GGGTGCCTCTGTCTCTCAAC |
| CTSS Forward | TCCTACCCTGGATCACCACT |
| CTSS Reverse | TTCTTCACTGGTCATGTCTCC |
| IL20RB Forward | GCTGATGCAACATCTGGGTTT |
| IL20RB Reverse | TGCATATGTTGGAGCTGAGG |
| LAT2 Forward | TTGCAACAGTTCTTGGAAACCC |
| LAT2 Reverse | GTTGCCTCTTGTGATGCGTG |
| IL18 Forward | AAGATGGCTGCTGAACCAGT |
| IL18 Reverse | GAGGCCGATTTCCTTGGTCA |
| OAS2 Forward | AGCTCTTTACTTTCCCCTTGGTT |
| OAS2 Reverse | GGAAACAGACAGGACGTGGA |
| PPIA Forward | CCCACCGTGTTCTTCGACATT |
| PPIA Reverse | GGACCCGTATGCTTTAGGATGA |
| HPRT1 Forward | CCTGGCGTCGTGATTAGTGAT |
| HPRT1 Reverse | AGACGTTCAGTCCTGTCCATAA |
| ACTB Forward | CTGTGGCATCCACGAAACTA |
| ACTB Reverse | CGCTCAGGAGGAGCAATG |
